# A conserved Pho4-Pho84 axis ubiquitously maintains intracellular phosphate homeostasis in yeast

**DOI:** 10.64898/2026.04.22.720069

**Authors:** Sreesa Sreedharan, Kshitija Shetty, Ganesh Muthu, Edrea Mendonca, Gaurav Singh, Felix Christian, Sunil Laxman

## Abstract

Inorganic phosphate (Pi) is central to fundamental cellular processes and the metabolic economy, and is constantly acquired in order to maintain intracellular Pi levels. While much is known about cellular adaptation during Pi starvation, how intracellular Pi is maintained in Pi-replete conditions remains unclear. Here, using *Saccharomyces cerevisiae*, we uncover an essential role for the Pho4 transcription factor in maintaining intracellular Pi under Pi-replete conditions, via the high affinity Pho84 transporter. Basal Pho4-dependent output is required for intracellular Pi maintenance, and the loss of Pho4 results in decreased intracellular Pi. We uncover that the Pho4 dependent, high affinity Pi transporter Pho84 is the primary transporter required for this intracellular Pi maintenance in phosphate replete conditions, and is not compensated by other transporters. The loss of Pho4 or Pho84 decreases intracellular Pi, with reduced ATP and glycolysis, and decreased growth. Through comparative genomic and phylogenetic analyses we establish that Pho84 is universally conserved across fungi, and Pho84 alone is orthologous to the plant high-affinity phosphate transporter PHT1. Thus, Pho84 is a primary determinant of intracellular Pi homeostasis during phosphate replete growth, and Pi acquisition in replete conditions is built around high-affinity phosphate transport. These findings reiterate the importance of Pi acquisition via high-affinity transport for metabolic homeostasis, with implications for microbial fermentation-based applications.

## Introduction

Phosphorus is an essential nutrient, and within cells, phosphate groups are widely distributed in ATP and other nucleotides, nucleic acids, phospholipids, phosphorylated proteins, and polyphosphate stores. Phosphorus is incorporated within cells in several organic and inorganic molecules, but is primarily acquired as inorganic phosphate (Pi). Pi is central to core cellular processes, and indispensable for overall homeostasis and the metabolic economy across all domains of life [1–3]. Pi is also directly consumed, released or coupled with many biochemical reactions, inhibits/activates key enzymes, and is critical for sustaining central carbon metabolic flux [4–7]. Cells therefore continuously require Pi to sustain their metabolism and growth, and therefore constantly acquire Pi from the external environment where it is typically abundant. However, two challenges exist for cells to meet this Pi demand: Pi is a charged molecule and does not freely diffuse across phospholipid membranes, and intracellular Pi levels must be sustained in proportion to ongoing metabolic demands for growth. Cells therefore have dedicated systems through which they continuously acquire Pi and maintain intracellular Pi. Surprisingly, intracellular Pi acquisition and maintenance in Pi-replete environments is taken for granted, and remains an unresolved question.

A lot of what we know about intracellular Pi homeostasis comes from studying responses to severe Pi starvation in model cells such as budding yeast (*Saccharomyces cerevisiae*), which activate well-studied adaptive programs to scavenge Pi and survive [8]. In budding yeast, which has been extensively used to identify responses and mechanisms in cells to Pi starvation, this is mediated by the PHO regulon, a transcriptional program activated during Pi limitation [9–12]. The Pho4 transcription factor controls these transcriptional programs, and is regulated by phosphorylation-dependent nucleo-cytoplasmic shuttling [13–16]: in Pi-replete conditions Pho4 is largely cytosolic, whereas under Pi limitation Pho4 accumulates in the nucleus and induces genes involved in Pi acquisition and mobilization, including Pi transporters and secreted phosphatases (Fig. 1A) [10,17,18]. This framework has defined our understanding of how cells respond to Pi starvation. However, whether this Pho4-PHO system contributes to intracellular Pi homeostasis during phosphate sufficiency is unclear.

**Figure 1:**
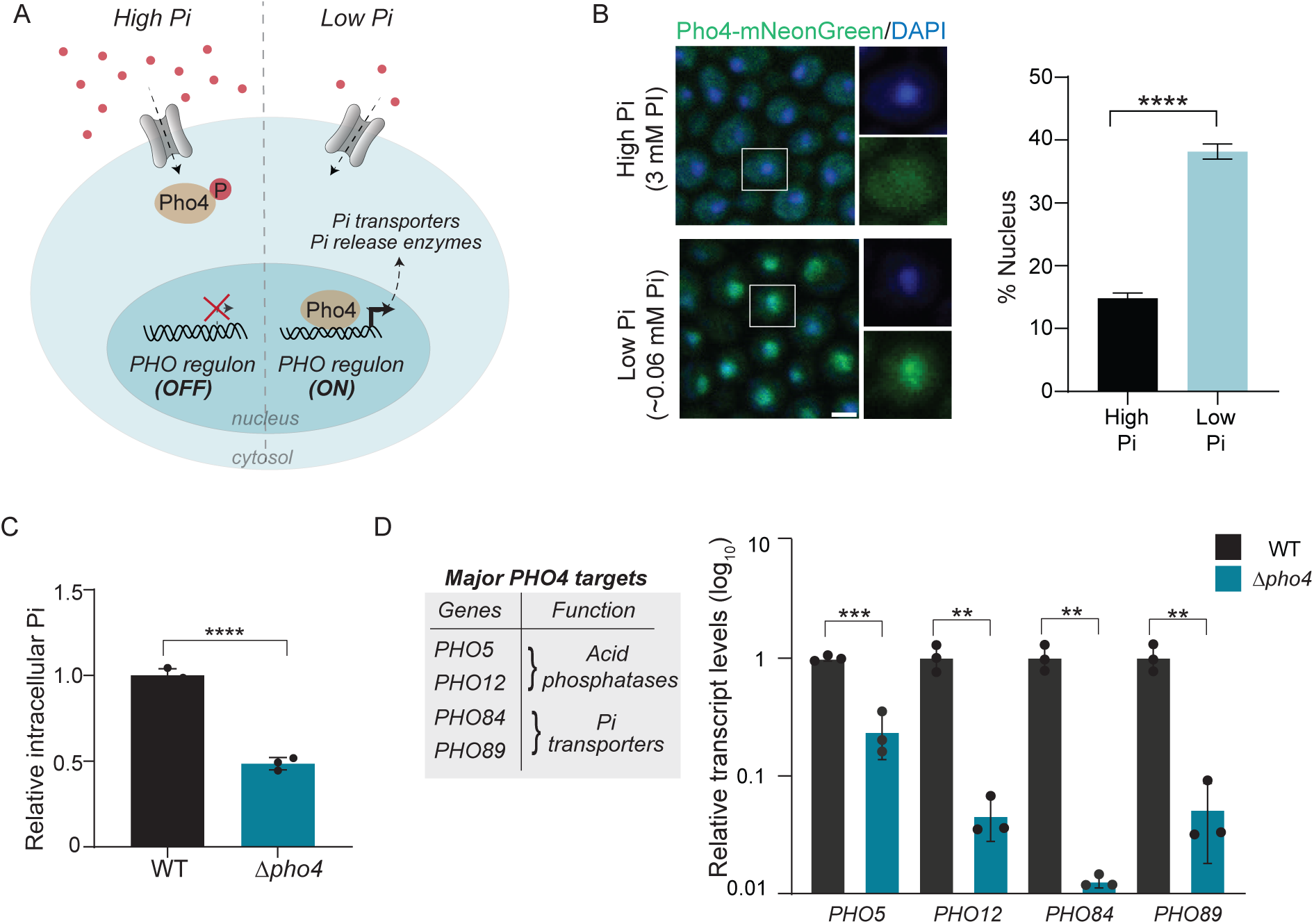
Pho4 is required to maintain intracellular Pi even in high Pi. **A. The Pho4-PHO regulon response system in *S.cerevisiae***. The schematic illustrates our current understanding of the Pho4-dependent PHO regulon in yeast. When extracellular Pi is high, Pho4 is phosphorylated and is predominantly cytoplasmic, keeping the PHO regulon repressed. Under low Pi, Pho4 accumulates in the nucleus and activates the PHO regulon/Pho4 responsive genes, which are involved in restoring Pi levels and managing starvation. **B. Subcellular localization of Pho4 and external Pi levels**. Left: Representative confocal images of Pho4–mNeonGreen in cells grown in high Pi (3 mM Pi) or low Pi (∼0.06 mM Pi) medium. Pho4 was tagged at its endogenous C-terminus with mNeonGreen. The nucleus was visualized using DAPI staining (see Materials and Methods for details). Scale bar = 2 µm. Right: quantification showing % nuclear localisation of Pho4 in the given conditions (see Materials and Methods for details). Also see Fig. S1A for Pi concentrations in the media. **C. Loss of *PHO4* (*Δpho4*) results in reduced intracellular Pi even in high Pi.** Relative intracellular Pi amounts in wild type (WT) and *Δpho4* cells in high Pi media (see Materials and Methods). Data are represented as mean ± SD from n = 3 biological replicates. Statistical significance was assessed using an unpaired two-tailed Student’s t-test (*Δpho4* vs WT): ****P<0.0001. **D. Pho4 dependent PHO target gene expression in high Pi.** Left: Summary table with four well-established Pho4 target genes, and their functions. Right: Relative transcript levels of these Pho4 target genes in WT and *Δpho4* cells in high Pi. Fold change values were calculated relative to WT cell gene expression, with *TAF10* as the reference gene. Data are represented as mean ± SD from n = 3 biological replicates. Statistical significance was calculated using unpaired Student’s t-tests (*Δpho4* vs WT): **P<0.01. ***P<0.001.

A primary way by which cells control intracellular Pi is through Pi uptake. Yeast encodes multiple Pi transporters that function under different extracellular Pi concentrations [19–22]. When Pi is abundant in the environment, uptake is generally assumed to depend on lower-affinity systems [23,24]. During Pi starvation, the Pho4 dependent, PHO-regulated high-affinity transporters such as Pho84 and Pho89 are strongly induced [17,25]. Because most studies have focused on starvation responses, the Pi acquisition architecture that maintains intracellular Pi during phosphate-replete growth has not been directly examined. However, this is important to understand because changes in phosphate uptake will affect cellular Pi pools, and subsequently carbon metabolism, energy balance, and growth rates, given the dependence of glycolysis and ATP turnover on Pi availability [4,5]. Understanding this therefore has implications on our understanding of basic cell growth-metabolism relationships, with important applications in biotechnology. For example, Pi uptake processes are important in microbial and yeast fermentation, secondary metabolite production, flavour production and many more [26,27]. Therefore, it is important to understand the phosphate transport systems that maintain intracellular Pi during Pi-replete growth, whether the Pho4/PHO-regulated transport contributes to this process, and how these constrain cell growth, biomass and metabolism. This question of how cells maintain Pi in replete conditions is also broadly relevant. Plants depend on Pi availability for their growth and development and relies on various strategies to acquire Pi from soil [28–31], in particular via the PHT1 family of transporters [32–34]. Thus, understanding how eukaryotic cells maintain intracellular Pi through defined uptake systems has broader relevance beyond fungal systems.

In this study, using *S. cerevisiae*, we identify an unexpected role for Pho4 in maintaining intracellular Pi in replete conditions. We find that Pho4- dependent, high-affinity phosphate transporter, Pho84, is the primary regulator of Pi intake, and is ubiquitously required to maintain intracellular Pi homeostasis during phosphate sufficiency. The loss of Pho4 or Pho84 alone phenocopy each other, and critically reduce intracellular Pi in cells. This intracellular Pi reduction due to the loss of Pho4 or Pho84 alters carbon metabolism, reduces glycolytic intermediates and ATP and decreases growth even when Pi is externally replete. Using comparative phylogenomic and protein analyses we find that the high-affinity Pho84 transporter is the primary Pi transporter present across the entire fungal kingdom, and orthologous to the critical PHT1 transporter system in plants. Together, these findings establish that the Pho4 dependent, Pho84 high-affinity phosphate transport system sustains Pi acquisition even during Pi sufficiency, and reveal how Pi homeostasis is built around high-affinity transport. These findings will impact engineering cell factories with optimized Pho84 amounts for fermentation based applications.

## Results

### Pho4 is required to maintain intracellular Pi even in high Pi

Cells obtain inorganic phosphate (Pi) exogenously, and Pi is central to cellular homeostasis. Pi is generally not limiting in media, and phosphate homeostasis responses have largely been studied during Pi limitation, especially using *S. cerevisiae* as a model eukaryotic cell [8,9,11,12]. The Pi starvation response in *S.cerevisiae* is mediated by the expression of the PHO regulon, a collection of transporters, phosphatases and enzymes that are involved in phosphate homeostasis. The expression of these genes are generally are thought to remain repressed when Pi is abundant externally (Fig. 1A). In yeast, their expression is regulated by the transcription factor Pho4, which is sequestered in the cytoplasm when Pi is abundant. During external Pi starvation, Pho4 localises to the nucleus, strongly activating the transcription of genes in this regulon (Fig. 1A). Given these observations with Pho4 localization and activity, the requirement of Pho4 for Pi homeostasis has been extensively studied only during Pi starvation [14–16,35], however, its requirement for phosphate homeostasis when Pi is replete is largely unknown. We therefore sought to first assess any role for Pho4 during homeostatic growth in Pi replete, standard medium (YPD, with ∼3 mM free Pi), and included controls of cells growing in low Pi (with ∼0.05-0.06 mM Pi) (Fig. S1 A). Note: all our experiments were carried out using a robust, fully prototrophic yeast strain from a CEN.PK background [36].

We first assessed the protein localization of the Pho4 transcription factor, in cells grown in either standard, Pi replete-media, or under Pi limitation. For this, we tagged *PHO4* at its endogenous C-terminus with a fluorescent tag, mNeonGreen. Cells were then grown in either high Pi media or low Pi media (Fig. S1 A). The subcellular localization of Pho4 was examined using confocal microscopy, and carefully quantified in these conditions. We observed that in high Pi, Pho4 was predominantly cytosolic as expected (Fig. 1B). Upon quantifying these images, however, we observe that ∼10-15% of the total cellular Pho4-mNeonGreen signal fell within the nuclear mask even in Pi replete media conditions (Fig. 1B). These data therefore indicates that a small population of Pho4 localises to the nucleus even in high Pi. Expectedly, in low Pi, Pho4 substantially accumulated in the nucleus, indicating its strong activation (Fig. 1B). These are consistent with known observations that establish canonical PHO pathway regulation, where Pho4 is localized within the nucleus upon Pi starvation.

Given this observation that there appears to be significant ‘basal’ nuclear Pho4 in high Pi, we asked if the Pho4 protein had any role in intracellular Pi maintenance when inorganic phosphate was not limiting in the medium, i.e. in standard medium. To address this, we grew wild type (WT) and *Δpho4* cells in standard, high Pi medium, and quantitatively estimated the amounts of intracellular Pi in these cells. Surprisingly, we found that *Δpho4* cells had a substantial, ∼50% reduction in intracellular Pi in these replete conditions (Fig. 1C). This suggests that Pho4 is required for cells to maintain intracellular Pi even in high Pi medium (Fig. 1C).

These data imply that the ‘basal’ Pho4-dependent gene expression is required to maintain intracellular Pi homeostasis. To assess this, we measured transcript levels of well-known Pho4 targets in WT and *Δpho4* cells grown in high Pi (Fig. 1D, left). These are PHO genes involved in Pi acquisition, including *PHO5* and *PHO12* (acid phosphatases) and *PHO84* and *PHO89* (Pi transporters) [17,18,37]. Transcript levels of all tested PHO targets were significantly reduced in *Δpho4* cells relative to WT (Fig. 1D, right). These results indicate that Pho4 dependent, PHO target gene expression persists in high Pi medium, suggesting that Pi acquisition even when inorganic phosphate is abundant requires Pho4 dependent activity, which is lost in *Δpho4* cells.

Collectively, this data reveal a necessary role of Pho4 in maintaining intracellular Pi, even when inorganic phosphate is externally abundant.

### Pho84 is the predominant Pho4 regulated Pi transporter in high Pi

Among the *PHO4* targets assessed, *PHO84*, (encoding a Pi transporter), showed the largest relative reduction in transcript levels in *Δpho4* cells (Fig. 1D). This raised the possibility that Pho84 (or other Pi transporters) could contribute to maintaining Pi homeostasis in high Pi conditions. However, currently the roles or hierarchies of different Pi transporters in Pi replete conditions are not established. To this end, we wondered if the decreased intracellular Pi in *Δpho4* cells was due to reduced Pi transporters and consequently Pi uptake. It has been generally assumed that Pi uptake in high Pi is attributed to constitutively present, low affinity Pi transporters: Pho87 and Pho90 [20,21,24], although their impact on actual, internal Pi is unknown. Further, the Pho4 dependent, high affinity transporters (Pho84 and Pho89) are not thought to have critical roles when Pi is abundant, and (as earlier described) have been primarily studied only during Pi starvation. Given our observations that Pho84 and Pho89 transcript levels in high Pi are Pho4 dependent, we first asked what is the relative abundance of the currently known Pi transporters, in cells growing in standard, high Pi media. To address this, we generated strains in which each of the Pi transporters were individually tagged at their endogenous C-termini with a FLAG epitope (Fig. 2A). We then measured their relative protein amounts, in high Pi by Western blotting (Fig. 2A, Fig. S2 A). Among all the transporters examined, Pho84 showed by far the highest protein amounts in Pi-replete conditions (Fig. 2A, Fig. S2 A). Pho89 (another high affinity Pi transporter) was present at much lower levels compared to Pho84 in the same condition. Finally, the lower-affinity transporters Pho87 and Pho90 were present at substantially lower levels than Pho84 in the same medium (Fig. 2A, Fig. S2 A). Thus, although Pho84 has been classically viewed as a starvation-induced Pi transporter, our data indicate that it is present at much higher levels than any other transporter in high Pi.

**Figure 2:**
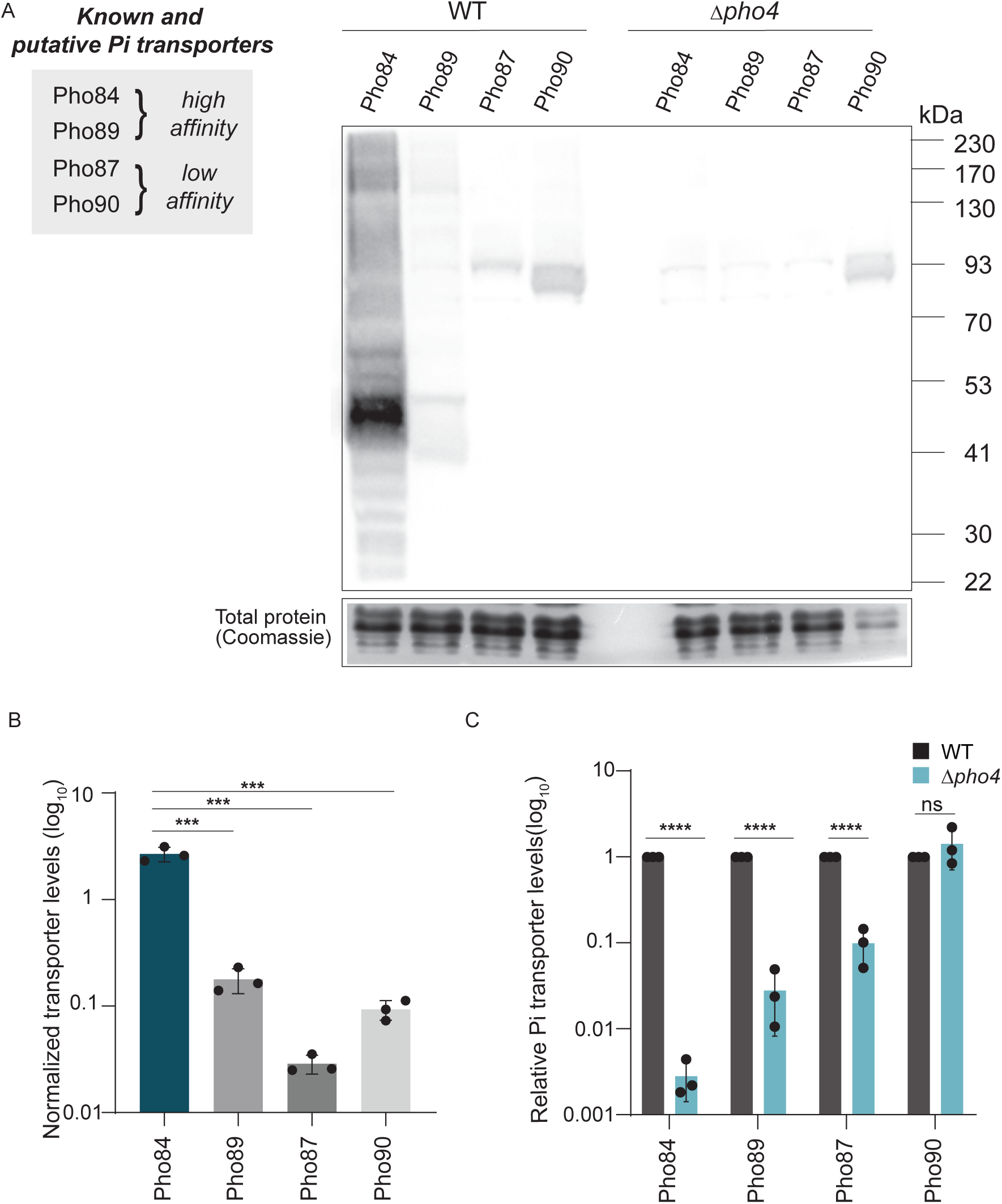
Pho84 is the predominant Pho4 regulated Pi transporter in high Pi. A. Relative amounts of known Pi transporters in high Pi in WT and *Δpho4* cells. Left: Summary of the major *S. cerevisiae* Pi transporters and their current classification as high- or low-affinity transporters. Right: Western blot analysis comparing levels of the indicated Pi transporters in WT and *Δpho4* cells in high Pi. Each transporter was chromosomally tagged at its respective endogenous C-terminus with a FLAG epitope and detected using an anti-FLAG antibody. Coomassie-stained gel portions were used as loading controls for total protein. A representative blot from n = 3 biological replicates is shown. Also see Fig. S2 A for a higher exposure blot B. Quantification of basal phosphate transporter abundance in WT cells grown in high Pi. Band intensities for the indicated FLAG-tagged transporters were quantified from western blots and normalized to the corresponding loading controls. Data shown are mean ± SD from n = 3 biological replicates. Statistical significance was calculated using unpaired two-tailed Student’s t-tests ***P<0.001 C. Relative amounts of all transporters in *Δpho4* cells, calculated and compared to the respective protein amount in WT cells (set as 1 for each protein). Data are represented as mean± SD from n = 3 biological replicates. Statistical significance was calculated using unpaired two-tailed Student’s t-tests ****P<0.0001; ns: non-significant difference

We next asked whether these transporter levels in high Pi depend on Pho4. To address this, we quantified and compared the relative abundance of the same transporters in WT and *Δpho4* cells grown in high Pi (Fig. 2A). We found that the abundance of all high affinity Pi transporters reduced in *Δpho4* cells, with the largest relative decrease seen for Pho84 (Fig. 2A, Fig. S2 A). Together, these results show that Pi uptake machinery depends on Pho4 even in high Pi. In particular, the strong reduction in Pho84 in *Δpho4* cells suggests that Pho84 may be important for cells to maintain intracellular Pi levels in Pi replete media.

### Pho84 maintains intracellular Pi in high Pi

Our data showed that the Pho84 Pi transporter is Pho4 dependent, and is present at high levels in high Pi. Since both low and high affinity transporters are present at varying abundances under these conditions, we asked which of them were required to maintain intracellular Pi levels. To address this, we measured intracellular Pi levels in individual mutants of these Pi transporters, in high Pi.

First, we observed that the loss of either of the low affinity transporters, Pho87 or Pho90, had no effect at all on intracellular Pi levels (as compared to WT cells), in high Pi (Fig. 3A). Next, the loss of Pho89 also did not significantly affect intracellular Pi levels (Fig. 3A). In contrast, the loss of Pho84 alone caused a marked reduction in intracellular Pi levels, and these levels were comparable to (or even lower) than *Δpho4* cells (Fig. 3A). These data establish that Pho84 is the primary transporter required to maintain intracellular Pi in phosphate replete media. The other transporters play a minor role in maintaining Pi levels in yeast. These results are also consistent with the abundance of each of the Pi transporters as seen in Fig. 2.

**Figure 3:**
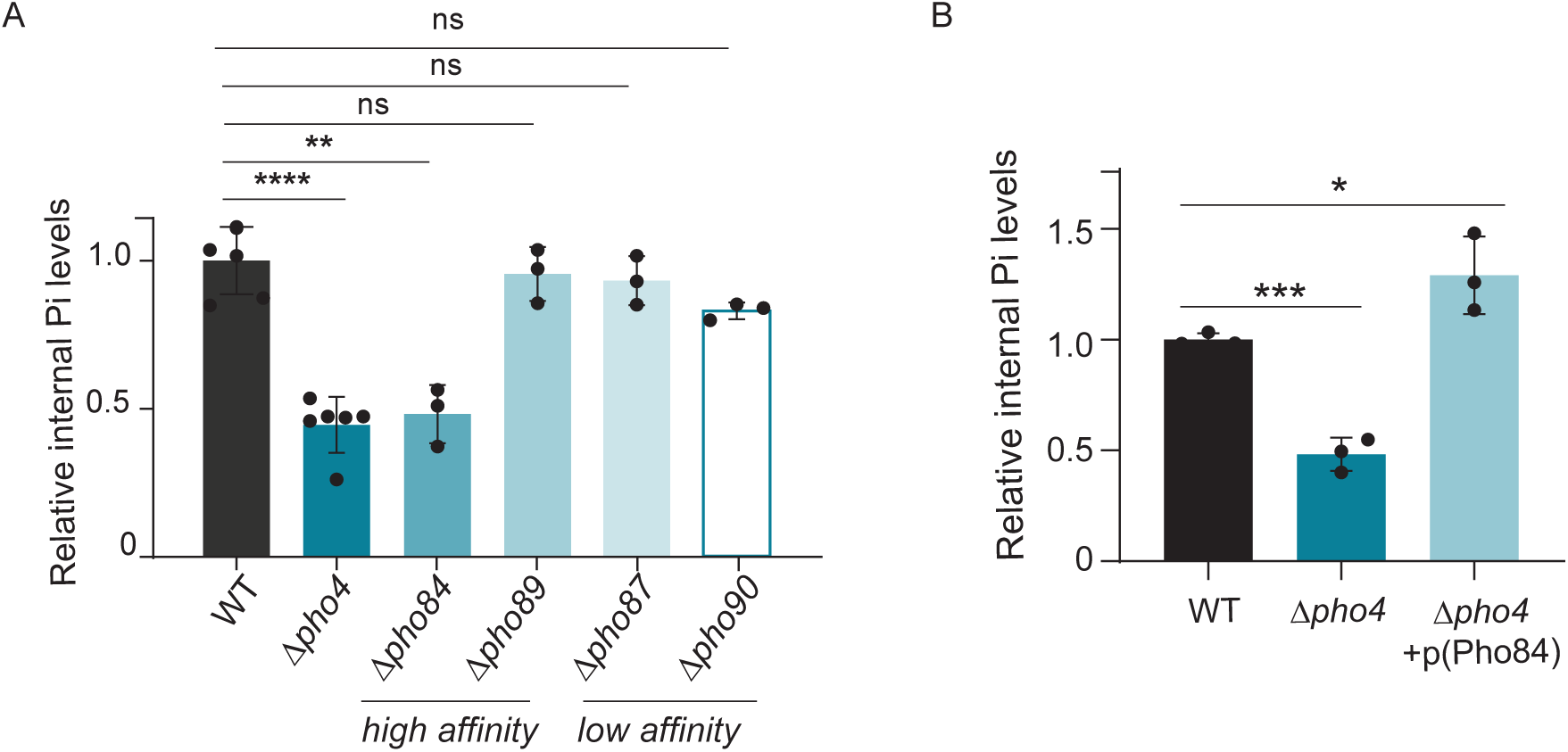
Pho84 maintains intracellular Pi in high Pi. **A. *PHO84* deletion alone phenocopies *Δpho4* in intracellular Pi levels.** Relative internal Pi levels in WT, *Δpho4* and in individual knockouts of each Pi transporter, from cells grown in high Pi. Data are represented as mean ± SD from n = 3 biological replicates. Statistical significance was assessed using an unpaired two-tailed Student’s t-test, comparing mutants to WT. **P<0.01, ****P<0.0001, ns: non-significant difference **B. Pho84 overexpression rescues intracellular Pi levels in *Δpho4* cells.** Relative intracellular Pi levels in WT, *Δpho4* and *Δpho4* cells with a plasmid overexpressing Pho84 (see Materials and Methods for details). The indicated strains were grown in high Pi media and intracellular Pi levels were measured. Data are represented as mean ± SD from n = 3 biological replicates. Statistical significance was assessed using an unpaired two-tailed Student’s t-test, comparisons to WT. *P<0.05, ***P<0.001. Also see Fig S3 A and S3 B

We next asked whether the intracellular Pi reduction in *Δpho4* cells was due to reduced Pho84. If this were so, then restoring Pho84 expression in *Δpho4* cells would rescue Pi levels. To test this, we overexpressed Pho84 (Pho84 OE) in *Δpho4* cells (using ectopic Pho84 expression plasmid under the control of a strong TEF1 promoter). The expression of Pho84 was confirmed by Western blot analysis (Fig. S3 A). We grew WT, *Δpho4*, and *Δpho4 +* Pho84 OE strains in high Pi and measured intracellular Pi levels. Notably, the overexpression of Pho84 in *Δpho4* increased and restored intracellular Pi levels (Fig. 3B). As expected, Pho84 OE also rescued *Δpho84* cells, and restored intracellular Pi levels in that background (Fig. S3 B). Together, these experiments reiterate that reduced Pho84 is causal to the intracellular Pi reduction observed in *Δpho4* cells.

### Pi depletion due to the loss of Pho84 is not compensated by other Pi transporters

Our data showed that Pho84 is abundant in high Pi, and that the loss of Pho84 alone reduces intracellular Pi levels under these conditions (Fig. 3A). In contrast, deletion of the other transporters did not measurably affect intracellular Pi (Fig. 3A). Since multiple Pi transport systems exist in yeast, one explanation for the lack of Pi reduction in these deletion strains could be functional compensation or redundancy, where the loss of one transporter might be compensated by another.

We therefore also assessed whether deletion of individual Pi transporters leads to compensatory changes in the abundance of other transporters, under high Pi. For this, we generated individual transporter knock-outs, where every other transporter was endogenously epitope-tagged, enabling comparison of transporter abundance across knockout backgrounds. We observed that the knock-outs of the low-affinity transporters Pho87 and Pho90, as well as the high-affinity transporter Pho89, led to a small increase in Pho84 alone (Fig. 4A, 4B). This was specific to Pho84, since we did not observe a similar increase in the levels of any other transporter (Fig. 4A, 4B). Thus, Pho84 is the only transporter that increases when alternative Pi transport routes were removed.

**Figure 4:**
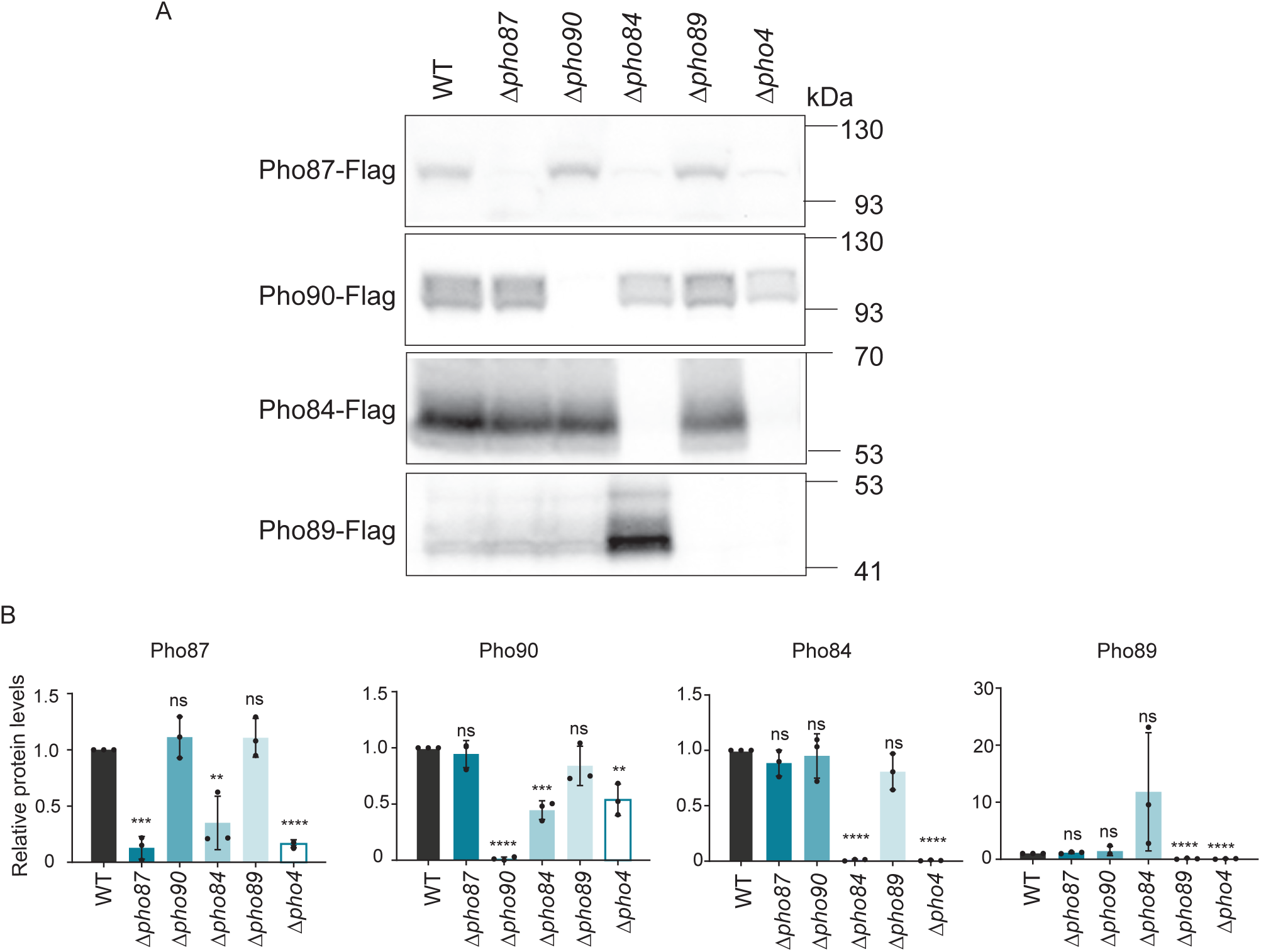
Pi depletion due to the loss of Pho84 is not compensated by other Pi transporters. **A.** Western blot analysis comparing levels of the major phosphate transporters in WT, *Δpho4*, and individual knockouts of Pi transporters grown in high Pi. In each knockout background, the indicated transporter(s) were tagged at the endogenous C-terminus with a FLAG epitope, and detected by anti-FLAG immunoblotting (see Materials and Methods for strain construction details). Note: for each transporter probed, its respective knockout strain served as the negative control (indicating absence of protein). Coomassie-stained gel portions were used as total protein loading controls. **B.** Graphs quantifying relative protein amounts for the indicated Pi transporters in WT, *Δpho4* and respective Pi transporter knockouts. Data are represented as mean+/−SD from n=3 biological replicates, all compared to WT. Statistical significance was calculated using unpaired Student’s t-test **P<0.01, ***P<0.001, ****P<0.0001, ns: non-significant difference

In contrast, in *Δpho84*, the levels of the other Pi transporters did not change sufficiently, to compensate for its loss (Fig. 4A, 4B). Thus, while the presence of Pho84 is sufficient to manage the loss of any other transporter, the converse is not true under high Pi, and the remaining transporters do not increase or compensate for the loss of Pho84.

Collectively, these results reiterate that Pho84 is the primary transporter required to maintain intracellular Pi levels in high Pi. They also explain why *Δpho84* uniquely reduces intracellular Pi: loss of Pho84 is not accompanied by compensatory increases in other transporters, whereas Pho84 itself is preferentially upregulated when other Pi transport routes are removed.

### Loss of Pho4 or Pho84 reduces glycolytic intermediates and ATP

Our results establish that both *Δpho4* and *Δpho84* cells have reduced intracellular Pi levels even in high Pi. Recent studies have shown that intracellular phosphate balance is linked to carbon metabolism, with changes in Pi availability being associated with altered glycolytic flux, trehalose synthesis, and mitochondrial activity [4,6,38,39]. This suggests that the Pi reduction in these strains could have broader consequences on glycolytic state, ATP levels, and eventually growth. We therefore next asked what the metabolic and growth consequences were in cells lacking Pho4 or Pho84, in high Pi.

To address this, we first measured glycolytic intermediates in WT, *Δpho4*, and *Δpho84* cells grown in high Pi, using highly targeted, quantitative LC/MS/MS approaches [40]. We observed that both *Δpho4* and *Δpho84* cells had significantly reduced levels of multiple glycolytic intermediates compared to WT (Fig. 5A). Thus, the intracellular Pi defect in these strains is accompanied by a reduced glycolytic signature.

**Figure 5:**
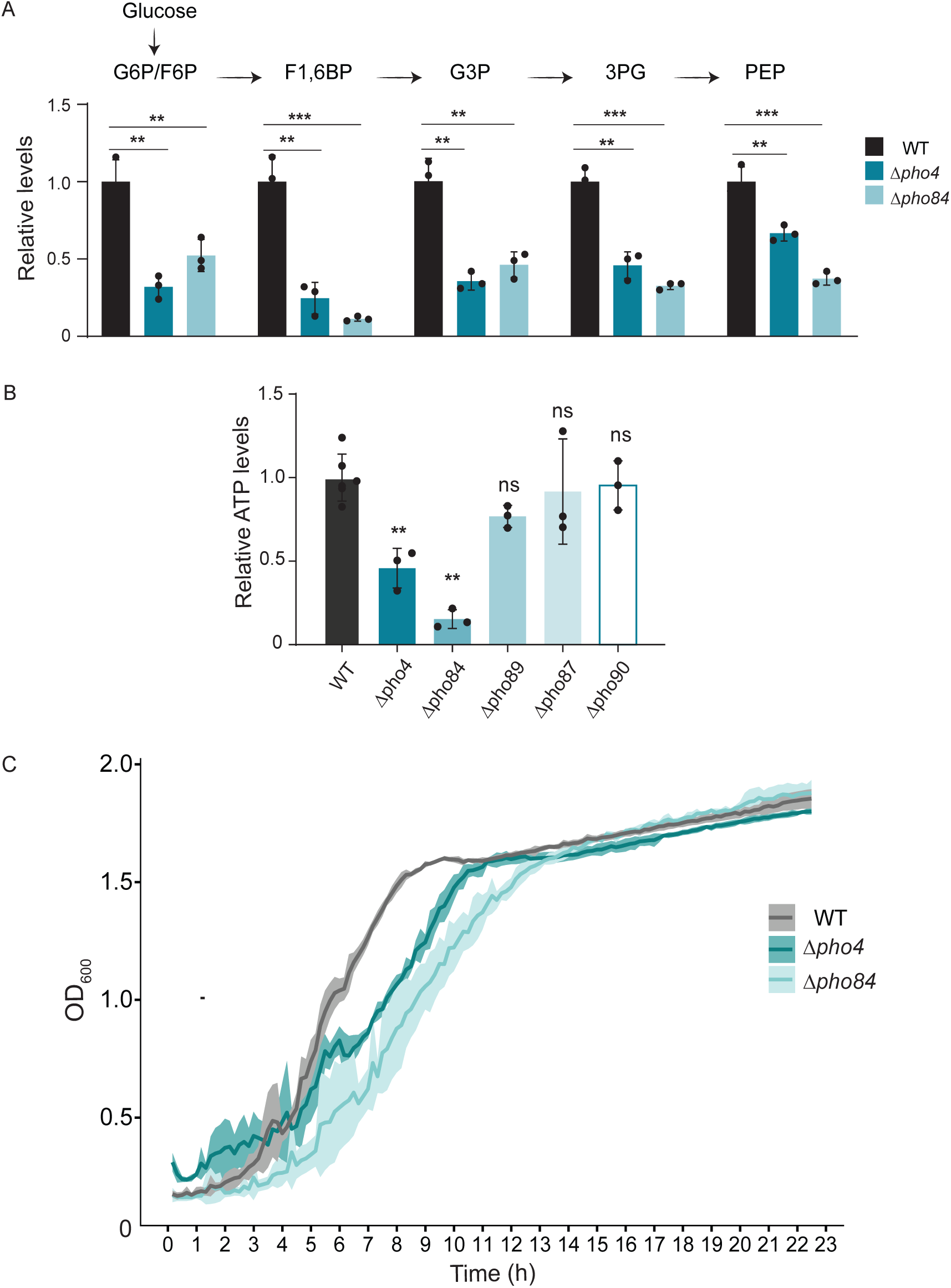
Loss of Pho4 or Pho84 reduces glycolytic intermediates and ATP. **A. Glycolytic intermediates decrease in *Δpho4* and *Δpho84* in high Pi.** Relative levels of glycolytic intermediates in WT, *Δpho4*, and *Δpho84* cells grown in high Pi. Metabolite levels were measured by LC-MS/MS analysis as described in Materials and Methods. Data are represented as mean+/−SD from n=3 biological replicates, all compared to WT. Statistical significance was calculated using unpaired Student’s t-test. **P<0.01, ***P<0.001. **B. *Δpho4* and *Δpho84* cells have reduced ATP in high Pi.** Cellular ATP levels in WT, *Δpho4*, and each individual Pi transporter knocked out, grown to log phase in high Pi medium, measured using luminescence based assay as described in Materials and Methods. Data are represented as meanv± SD from n = 3 biological replicates, comparing mutants to WT. Statistical significance was determined using unpaired Student’s t-tests. **P<0.01, ns: non-significant difference. **C. *Δpho4* and *Δpho84* cells have decreased growth in high Pi.** Growth curves of WT, *Δpho4*, and *Δpho84* cells in high Pi medium. Cultures were inoculated at OD_600_ = 0.2, and OD_600_ was measured every 10 minutes for ∼24 h. *Δpho4* and *Δpho84* cells displayed reduced log phase growth rate under these conditions. Data are represented as mean ± SD (solid lines = mean; shaded region = SD) from n=3 biological replicates.

We next asked whether this is also reflected in ATP levels. To test this, we measured ATP in WT, *Δpho4*, and *Δpho84* cells grown under the same conditions of high Pi. Consistent with the decrease in glycolytic intermediates, ATP levels were substantially reduced in both *Δpho4* and *Δpho84* cells relative to WT (Fig. 5B). These data indicate that loss of Pho4 or Pho84 is associated with a lower energetic state even in high Pi.

Finally, we asked whether these metabolic defects in *Δpho4*, and *Δpho84* cells are reflected in growth. We therefore compared the growth of WT, *Δpho4*, and *Δpho84* cells in high Pi over time (Fig. 5C). Both knock-out strains showed (comparable) reduced growth compared to WT under these conditions. Thus, the intracellular Pi defect in *Δpho4* and *Δpho84* cells is accompanied by both metabolic and growth defects even in Pi-replete medium.

Collectively, these results establish that the reduced intracellular Pi phenotype of *Δpho4* and *Δpho84* cells is not restricted to Pi levels alone, but is associated with altered metabolic state, ATP stress and impaired growth in standard media environments where Pi is not limiting.

### Pho84 is broadly conserved across fungi and shares orthology with plant PHT1 transporters

Our results in *S. cerevisiae* indicate that Pho84 is the primary phosphate transporter required for intracellular Pi maintenance even in high Pi medium. We therefore wondered whether the transporter architecture highlighted by our yeast data is conserved more broadly across fungi. To address this, we performed a comparative phylogenomic analysis of phosphate transporters, following our previously described phylogenetic methodology [41] across 193 fungal genomes, spanning Ascomycota (149), Basidiomycota (36), and several early-diverging fungal lineages (Fig. 6A).

**Figure 6:**
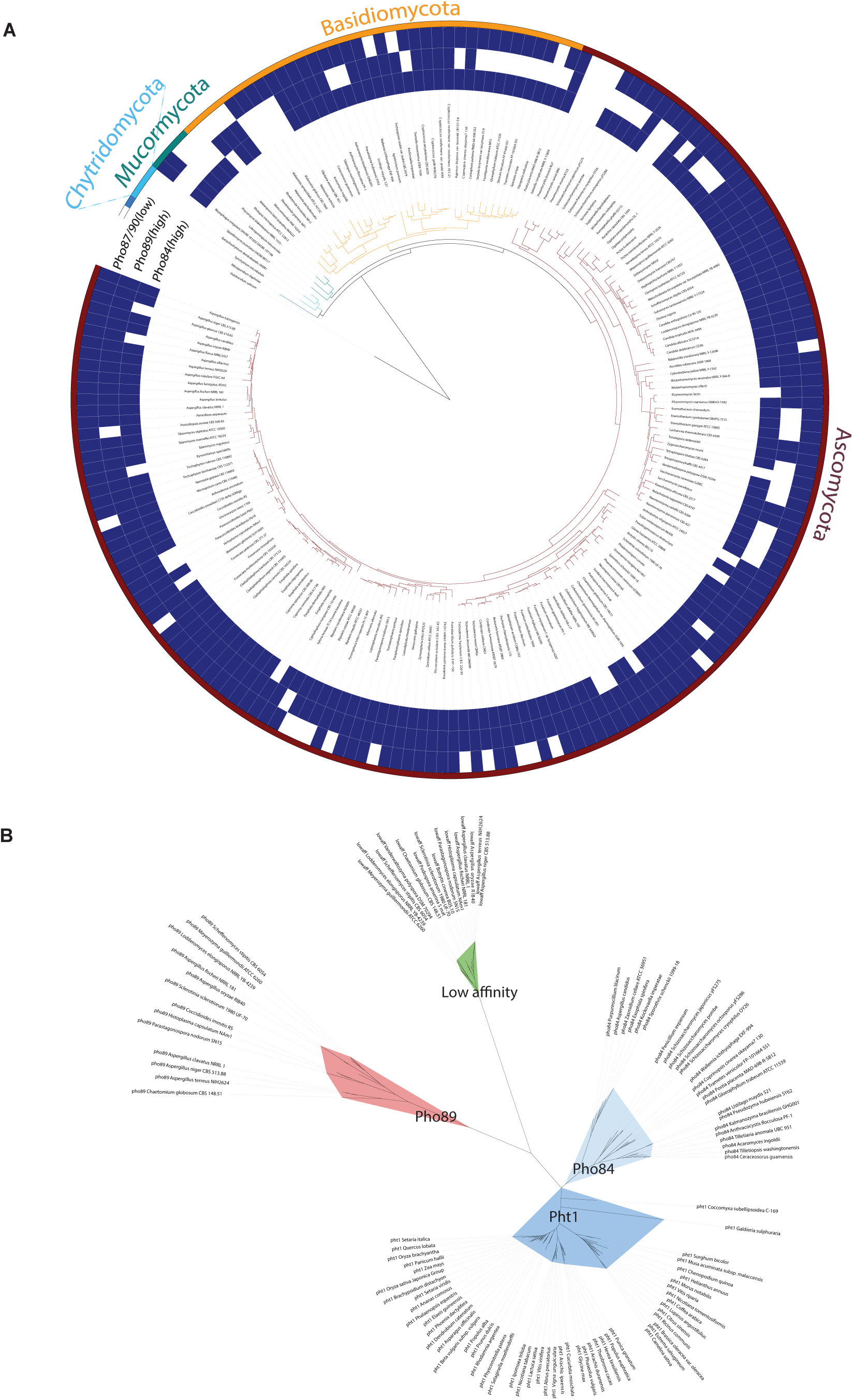
Pho84 is broadly conserved across fungi and shares orthology with plant PHT1 transporters. A. Comparative genomics analysis of phosphate transporters across 193 fungal genomes spanning Ascomycota, Basidiomycota, Mucormycota and Chytridomycota species. The fungal species tree shown was constructed using 27 orthologous genes (more details in the Materials and Methods section). The outer panels in the phylogenetic tree show heatmaps indicating the presence (blue) or absence (white) of the phosphate transporters across species. High-affinity transporters are represented separately as Pho84 and Pho89. Low-affinity transporters (Pho87, Pho90) were grouped into a single track for clarity, since they share the same core protein domains and belong to the same orthologous group. B. Maximum likelihood phylogenetic tree of eukaryotic phosphate transporters. The gene tree illustrates the evolutionary relationships among fungal high-affinity (PHO84, PHO89), fungal low-affinity, and plant (PHT1) phosphate transporter proteins. Plant PHT1 and fungal PHO84 form a closely unified clade (highlighted in blue shades) indicating an orthologous relationship. In contrast, fungal PHO89 (highlighted in red) and the fungal low-affinity transporters (highlighted in green) diverge into entirely separate, distinct lineages.

We first asked how widely the different phosphate transporters are distributed across the fungal kingdom. We found that the phosphate transporters are nearly ubiquitous in fungi, being present in 96.3% of the fungal genomes examined (Fig. 6A). Thus, dedicated phosphate uptake systems are broadly conserved across fungi. Next, we compared the distribution of high and low-affinity phosphate transporters. High-affinity transporters (Pho84 and Pho89) are prevalent and are present in 92.7% of the fungi analysed. Of these, the Pho84 transporter is by far the most prevalent transporter, being present in 85.5% of fungal species. Pho89 is present only in 72.5%. In addition, 65.3% of species encoded both Pho84 and Pho89, indicating that dual high-affinity transport systems are moderately widespread across fungi (Fig. 6A).

Since Pho87 and Pho90 share the same core protein domains (PF03600- citrate transporter; PF03105- SPX domain) and belong to the same orthologous group (K14430, phosphate transporter), we grouped these proteins into a single low-affinity transporter category for clarity in the phylogenetic analysis. We find that the low-affinity transporters together were broadly represented and detected in 84.4% of the fungi analysed (Fig. 6A).

Together, these results show that phosphate transporters are broadly conserved across fungi, with high-affinity transport being especially prevalent. In particular, the widespread conservation of Pho84 suggested that this transporter may represent a central and conserved component of fungal phosphate uptake architecture.

We next asked whether the strong conservation of Pho84 across fungi extends beyond the fungal kingdom. We asked whether any of the fungal phosphate transporters belong to a more widely conserved phosphate uptake module. In this context, plant growth and development are critically coupled to the availability of Pi, where they utilise transport systems to uptake Pi from the soil [29,31,42]. The major plasma membrane Pi uptake system in plants is the PHT1 family of transporters [33,34]. We therefore first compared all the fungal phosphate transporters with the well-studied plant phosphate transporter PHT1 (Fig. 6B). Notably, among all the fungal transporters examined, only Pho84 showed clear similarity to PHT1, grouping closely together (Fig. 6B). The Pho84 and PHT1 belong to the same orthologous group (K08176; MFS transporter, PHS family, inorganic phosphate transporter). In contrast, the Pho89 transporter, and the low-affinity transporters did not, and clearly fell as outgroups from the plant PHT1 transporter. Thus, Pho84 is part of a more widely conserved, high-affinity phosphate transport module that extends to plants.

Collectively, these results reveal Pho84 as a most widely conserved high-affinity phosphate transporter across fungi, and place it within a broad, conserved phosphate uptake module extending beyond the fungal kingdom.

## Discussion

In this study, using *S. cerevisiae* we identify an essential role for the Pho4-PHO system in maintaining intracellular Pi homeostasis for Pi-replete growth. “Basal” Pho4 is required to maintain intracellular Pi levels and homeostasis, via its downstream targets (Fig. 1). In particular, the Pho4 dependent high affinity Pi transporter, Pho84 emerges as the major Pi transporter required to maintain intracellular Pi: it is the most abundant transporter in Pi replete conditions, its loss alone results in reduced intracellular Pi (Fig. 2 and 3). Notably, Pho84 loss is not compensated by other transporters, further reiterating a critical role for Pho84 in maintaining intracellular Pi in inorganic phosphate replete environments (Fig. 4). The loss of either Pho4 or Pho84, and the resultant reduction in internal Pi, results in cells exhibiting clear signatures of ‘starvation’ including reduced glycolytic intermediates, ATP, and slower growth (Fig. 5). Together, these results identify Pho4-dependent Pho84 as a major determinant of intracellular Pi homeostasis during Pi-replete growth.

Inorganic phosphates are crucial for cellular homeostasis across different domains of life [1,43,44]. For all cells, they are indispensable for metabolism and energetics and forms a central component of phospholipid membranes, nucleic acids, nucleotides etc. [5,6,8]. For plants and other organisms, they are critical for growth and development [30,42]. Therefore, it is not surprising that organisms have devised multiple strategies to maintain sufficient intracellular Pi concentrations, to sustain growth. Most work on Pi homeostasis in plants, bacteria and yeast have focused on transcriptional responses to extreme Pi starvation [45–47]. In yeasts *(S.cerevisiae)*, this is mediated by the PHO regulon where Pho4 enters the nucleus and induces the PHO regulon, encoding genes involved in Pi acquisition or recycling [9,11,12,14,16]. This framework has defined how cells manage Pi limitation.

Contrastingly, less attention has been given to how cells continuously maintain intracellular Pi during routine growth, despite knowledge that Pi demand is high [4,5,48]. This implies that cells have efficient Pi transport systems that will continuously import Pi. While the low affinity transport system have been presumed to support Pi uptake in high Pi environments, this assumption has not been systematically assessed. Our study reveals that the cells have built Pi acquisition for their economies using high affinity transporter systems. Pho84 is by far the most predominant Pi transporter, and its expression is critical to maintain internal Pi levels and growth. Notably, this high affinity Pi transporter is near-ubiquitously conserved across the entire fungal kingdom (Fig. 6), making Pho84 the predominant Pi transporter in all fungi. Additionally, all plants rely critically on the high-affinity Pht1 transporter for their (considerable) Pi requirements. The Pht1 transporter and the Pho84 fungal transporter are highly conserved, and belong to the same family of transporters (Fig. 6), while other Pi transporters are clear outgroups. These findings reveal that a plurality of eukaryotes have evolved their intracellular phosphate economies around high affinity Pi transporter systems.

The dependence on high affinity transporters for Pi uptake in replete conditions is seemingly surprising, especially because these are known to be induced in low µM concentrations of Pi ([20]. However, building Pi acquisition around high affinity Pi transporters, regardless of external Pi concentration, is in fact logical. Even though Pi is abundant, transient fluctuations in external availability or a constant increase in cellular demands that exceeds availability (for example during cell division, metabolism etc.) ([8,48,49], necessitates the need for better efficient transport systems, that can ensure a continuous supply of Pi. In the case of plants, though phosphorous is abundant, the amount of bioavailable inorganic phosphates (preferred in plants) is low. This is because in soil they are usually chelated to metal ions and have to be pumped up a concentration gradient [34,42,50]. Therefore, across systems, the use of a high-affinity transporter in high Pi can provide a robust way for cells to maintain high intracellular Pi even in all conditions. Interestingly, having Pi acquisition built around high affinity transporter could be useful not only for Pi uptake, but also for extracellular Pi sensing and overall nutrient signalling. Indeed, recent studies suggest that Pho84 functions as a Pi ‘transceptor’, coupling Pi binding to signalling in addition to transport [51], and our systems-level data now substantiates this possibility.

Further, the ‘basal’ activity requirement of Pho4 is very poorly understood, given the contrasting, strong activation observed during Pi starvation. Our data firmly establishes that the basal expression of a subset of PHO genes, particularly the high affinity Pho84 transporter, is critical for Pi maintenance. Our findings therefore focus on the importance of Pi acquisition itself, and strikingly compliment recent studies proposing a ‘transporter centric’ role for Pho4 [52]. Our studies also reiterate early, mechanistic studies showing that partially phosphorylated Pho4 can localise to the nucleus and activates only a subset of Pi-responsive genes in certain contexts [52,53]).

Our results intriguingly raise the question of why cells might have both low and high-affinity Pi transporter systems. The simplest view is that these systems operate in a strict starvation-versus-replete context [20,54,55]. Our data suggest that this is likely too simple an assumption. Pho84 is clearly the main determinant of intracellular Pi levels, whereas the low-affinity transporters do not have the same effect on steady-state intracellular Pi. At the same time, this does not mean that the low-affinity systems are unimportant. They can function at other external Pi concentrations, during transitions in Pi availability, or under other physiological states not captured in our assays, or be coupled to signaling systems. In this regard, the compensatory increase in Pho84 when other transporters are removed, together with the lack of a comparable increase in other transporters upon Pho84 deletion (Figure 4), reiterates that Pho84 has a distinct and non-redundant function. Our data also indicate that not all high-affinity transporters contribute equally: although Pho89 is classically grouped with Pho84 as a high-affinity transporter, it is present at lower levels in high Pi medium and does not show the same effect on intracellular Pi. Thus, in high Pi, the uptake system appears to be organized around Pho84. More broadly, our data highlights why the role of the PHO system in high Pi conditions has remained underappreciated. Previous work has largely examined PHO output mainly through the lens of starvation induction, and not in contexts when external Pi is sufficient.

Finally, our data suggest that this replete-state Pi homeostasis, built around basal Pho4 activity, and the high affinity Pho84 transporter availability, has critical physiological consequences. Although textbook knowledge, it is often underappreciated that intracellular Pi balance is closely tied to central carbon metabolism and metabolic homeostasis [4,5,38,39]. The altered glycolytic state, reduced ATP levels, and impaired growth associated with the loss of (either) *PHO4* or *PHO84* reveal that maintaining intracellular Pi is not merely about preventing Pi depletion, but critically supports the metabolic state required to sustain high growth. Pho84-dependent Pi transport becomes critical for cells to match Pi supply to the metabolic flux, observed in standard, Pi-replete medium. The high affinity Pho84 transporter sustains Pi entry, and stabilizes intracellular Pi against changing metabolic demand. Finally, our comparative genomic analysis shows a broad conservation of Pho84. This becomes relevant for widely utilized industrial and fermentation settings with fungi, where growth robustness, biomass and metabolic outputs are critical parameters for bioproduction. Managing and engineering high affinity Pi transporter expression can therefore be used to constrain precision fermentation outputs. Taken together, our findings reveal that the Pho4-regulated, high-affinity Pi transport system of Pho84 is central to intracellular Pi homeostasis, sustaining high metabolism and growth during Pi-replete conditions.

## Materials and Methods

### Yeast strains, media and growth conditions

All strains used in this study are listed in table 1. Prototrophic, CEN.PK, haploid, mat ‘a’ mating type of *S.cerevisiae* was used in all experiments (Van Dijken et al., 2000).Gene deletions and chromosomal C-terminal epitope tagging was done as described elsewhere (Longtine et al., 1998).High Pi media used throughout the study was YPD (1% yeast extract, 2% peptone and 2% glucose) media. For solid media, granulated agar was added to a final concentration of 2%. Low Pi media was prepared as described in (Vengayil et al., 2024).For all experiments, the required strains were grown overnight in high Pi media at 30°C, 250 rpm. This primary culture was used to start a secondary culture at ∼OD600 = 0.2. Cells were allowed to grow to log phase (OD600 = ∼0.8-1) for all experiments.

**Table 1:** List of strains.

| S.No | Strain Name | Genotype | Description | Source |
| --- | --- | --- | --- | --- |
| 1 | CEN.PK a (WT) | Mat a | Wild type, haploid strain of CEN.PK with mat a mating type | [36] |
| 2 | Pho4-mNeonGreen | CEN.PK a *PHO4-mNeonGreen::kanMX6* | C terminal tagged Pho4 with mNeonGreen | This study |
| 3 | *Δpho4* | CEN.PK a *Δpho4::hphMX6* | Deletion of *PHO4* | This study |
| 4 | *Δpho84* | CEN.PK a *Δpho84::hphMX6* | Deletion of *PHO84* | This study |
| 5 | *Δpho89* | CEN.PK a *Δpho89::natMX6* | Deletion of *PHO89* | This study |
| 6 | *Δpho87* | CEN.PK a *Δpho87::hphMX6* | Deletion of *PHO87* | This study |
| 7 | *Δpho90* | CEN.PK a *Δpho90::natMX6* | Deletion of *PHO90* | This study |
| 8 | Pho84-FLAG | CEN.PK a *PHO84-3xFLAG::hphMX6* | C terminal tagged Pho84 | This study |
| 9 | Pho89-FLAG | CEN.PK a *PHO89-3xFLAG::natMX6* | C terminal tagged Pho89 | This study |
| 10 | Pho87-FLAG | CEN.PK a *PHO87-3xFLAG::kanMX6* | C terminal tagged Pho87 | This study |
| 11 | Pho90-FLAG | CEN.PK a *PHO90-3xFLAG::hphMX6* | C terminal tagged Pho90 | This study |
| 12 | Pho84-FLAG *Δpho4* | CEN.PK a *PHO84-3xFLAG::hphMX6 Δpho4::kanMX6* | C terminal tagged Pho84 in a *PHO4* deletion strain | This study |
| 13 | Pho89-FLAG *Δpho4* | CEN.PK a *PHO89-3xFLAG::natMX6 Δpho4::hphMX6* | C terminal tagged Pho89 in a *PHO4* deletion strain | This study |
| 14 | Pho87-FLAG *Δpho4* | CEN.PK a *PHO87-3xFLAG::kanMX6 Δpho4::hphMX6* | C terminal tagged Pho87 in a *PHO4* deletion strain | This study |
| 15 | Pho90-FLAG *Δpho4* | CEN.PK a *PHO90-3xFLAG::hphMX6 Δpho4::kanMX6* | C terminal tagged Pho90 in a *PHO4* deletion strain | This study |
| 17 | Pho84-FLAG *Δpho89* | CEN.PK a *PHO84-3xFLAG::hphMX6 Δpho89::natMX6* | C terminal tagged Pho84 in a *PHO89* deletion strain | This study |
| 18 | Pho84-FLAG *Δpho87* | CEN.PK a *PHO84-3xFLAG::hphMX6 Δpho87::natMX6* | C terminal tagged Pho84 in a *PHO87* deletion strain | This study |
| 19 | Pho84-FLAG *Δpho90* | CEN.PK a *PHO84-3xFLAG::hphMX6 Δpho90::natMX6* | C terminal tagged Pho84 in a *PHO90* deletion strain | This study |
| 20 | Pho89-FLAG *Δpho84* | CEN.PK a *PHO89-3xFLAG::natMX6 Δpho84::hphMX6* | C terminal tagged Pho89 in a *PHO84* deletion strain | This study |
| 22 | Pho89-FLAG<br>*Δpho87* | CEN.PK a *PHO89-3x*<br>*FLAG::natMX6*<br>*Δpho87::hphMX6* | C terminal tagged<br>Pho89 in a *PHO87*<br>deletion strain | This study |
| 23 | Pho89-FLAG<br>*Δpho90* | CEN.PK a *PHO89-3x*<br>*FLAG::natMX6*<br>*Δpho90::hphMX6* | C terminal tagged<br>Pho89 in a *PHO90*<br>deletion strain | This study |
| 24 | Pho87-FLAG<br>*Δpho84* | CEN.PK a *PHO87-3x*<br>*FLAG::kanMX6*<br>*Δpho84::natMX6* | C terminal tagged<br>Pho87 in a *PHO84*<br>deletion strain | This study |
| 25 | Pho87-FLAG<br>*Δpho89* | CEN.PK a *PHO87-3x*<br>*FLAG::kanMX6*<br>*Δpho89::natMX6* | C terminal tagged<br>Pho87 in a *PHO89*<br>deletion strain | This study |
| 27 | Pho87-FLAG<br>*Δpho90* | CEN.PK a *PHO87-3x*<br>*FLAG::kanMX6*<br>*Δpho90::natMX6* | C terminal tagged<br>Pho87 in a *PHO90*<br>deletion strain | This study |
| 28 | Pho90-FLAG<br>*Δpho84* | CEN.PK a *PHO90-3x*<br>*FLAG::hphMX6*<br>*Δpho84::natMX6* | C terminal tagged<br>Pho90 in a *PHO84*<br>deletion strain | This study |
| 29 | Pho90-FLAG<br>*Δpho89* | CEN.PK a *PHO90-3x*<br>*FLAG::hphMX6*<br>*Δpho89::natMX6* | C terminal tagged<br>Pho90 in a *PHO89*<br>deletion strain | This study |
| 30 | Pho90-FLAG<br>*Δpho87* | CEN.PK a *PHO90-3x*<br>*FLAG::hphMX6*<br>*Δpho87::natMX6* | C terminal tagged<br>Pho90 in a *PHO87*<br>deletion strain | This study |
| 32 | *Δpho4+* Pho84-<br>FLAG OE | CEN.PK a<br>*Δpho4::hphMX6* +<br>*pTEF1-PHO84-*<br>*FLAG::kanMX6* | Pho84-Flag<br>overexpression under<br>constitutive TEF1<br>promotor in *PHO4*<br>deletion strain | This study |
| 33 | *Δpho84+* Pho84-<br>FLAG OE | CEN.PK a<br>*Δpho84::hphMX6* +<br>*pTEF1-PHO84-*<br>*FLAG::kanMX6* | Pho84-Flag<br>overexpression under<br>constitutive TEF1<br>promotor in *PHO84*<br>deletion strain | This study |

### Confocal microscopy for Pho4 localization

#### Sample preparation for imaging

Cells with Pho4 tagged at the C terminus with a fluorescent mNeonGreen tag was grown in high Pi media to log phase. A portion of this culture was shifted to low Pi media for 2 hours. Samples corresponding to ∼2-3 OD600 cells were collected from both high Pi and low Pi grown cultures and fixed using paraformaldehyde at a final concentration of 4% for 20 minutes at room temperature (RT). The cells were washed thrice with 1X PBS (1000g, 2 minutes each). Cells were then permeabilized using 0.1% Triton X-100 for 5 minutes and washed again in 1X PBS thrice. Samples were incubated with DAPI (final concentration: 1ug/mL) for 5 minutes at RT followed by washing thrice in 1X PBS. Samples were resuspended in 1X PBS, mounted on slides and visualised.

#### Image acquisition and analysis

All fluorescence imaging experiments were performed on fixed cells using an Olympus FV3000 inverted confocal laser scanning microscope. A 60× oil-immersion objective (NA ∼1.4) was used, and fluorescence was detected using GaAsP-PMT high-sensitivity detectors. Pho4-tagged mNeonGreen was excited with a 488 nm laser, and nuclei stained with DAPI was excited with a 405 nm laser. For each condition, 3D confocal stacks were acquired in both channels, and each field of view contained a large population of cells.

Images were analyzed in ImageJ/Fiji using a custom written macro. For each z-stack image, the DAPI channel (channel 1) was used for nuclear segmentation and the mNeonGreen channel (channel 2) was used for signal quantification and estimation of whole-cell area. Nuclei were segmented from the projected channel 1 image using 3-cluster k-means clustering. The cluster with the highest mean channel 1 intensity was selected as the nuclear class, converted to a binary mask, despeckled, watershed-separated, and size-filtered to retain nuclei within the specified area range. A cell mask was generated from the projected channel 2 image by converting to 8-bit, enhancing contrast, applying the Default auto-threshold, converting to a binary mask, despeckling, and filling holes. The final cell quantification mask was defined as the union of the channel 2 cell mask and the filtered nuclear mask. The percentage of channel 2 signal in the nucleus was calculated as the integrated channel 2 intensity within the nuclear mask divided by the integrated channel 2 intensity within the final cell mask, multiplied by 100.

### Inorganic phosphate (Pi) measurement

Pi measurement was done using a malachite green (MG) phosphate assay kit (Cayman Chemicals, 10009325) as described in [39]. Briefly, WT or mutant cells were grown to log phase in high Pi media (OD600 of ∼0.8–1.0), and equal OD600 cells were collected by centrifugation (1000 × g, 2 min, RT). Cells were washed once with MS grade water (to prevent Pi contamination) (1000xg, 30 sec) and the pellets were flash frozen. For the assay, the pellets were first resuspended in 1 mL MS grade water and lysed by bead-beating (twice for 30 s, with 2 min incubation on ice in between). The lysate was cleared by centrifugation (14000 rpm, 10 min, 4°C). Protein amounts in the lysate were estimated by Bicinchoninic acid assay (BCA assay kit, G-Biosciences). Lysates were normalised for equal amounts of protein and the Pi amounts in the lysates were measured using the assay kit following manufacturer’s instructions. For extracellular Pi measurement, 1 mL of high and low Pi was used and diluted with MS grade water. Absolute Pi amounts in the media were calculated using the standard curve in the malachite green assay.

### Protein extraction and western blot analysis

Required strains were grown as described and ∼10 OD600 cells were collected by centrifugation, washed once and stored until further processing. Protein extraction and western blot analysis was done as described in detail in [57]. Briefly, cells were lysed by bead beating using 10% ice-cold trichloroacetic acid (TCA). Lysates were cleared by centrifugation, proteins precipitated and resuspended in SDS-glycerol buffer. Protein amounts in the samples were measured using BCA assay kit. Samples were normalised for equal protein, and was resolved on 4-12% bis tris gels (NUPAGE NP0322BOX). Relevant portions of the gel were cut and transferred to a nitrocellulose membrane and the bottom portion of the gel was stained with Coomassie Brilliant Blue for total protein normalization. Blots were blocked with 5% non-fat skim milk for 20 minutes. Then these were probed with anti-FLAG antibody raised in mouse (1:2000; Sigma-Aldrich, F1804) and anti-mouse horseradish peroxidase-conjugated secondary antibody (1:4000; Cell Signaling Technology, 7074S). The blot was developed using a standard enhanced chemiluminescence reagent (WesternBright ECL HRP substrate, Advansta K12045). Quantification of the blots and loading controls were performed using ImageJ, and statistical significance was determined using Student’s *t*-test in GraphPad.

### RNA extraction and quantitative RT-PCR

Cells were grown in high Pi media to log phase and equal OD600 cells were collected by centrifugation at 1000g, 2 minutes, 4°C. Total RNA extraction was carried out using the hot acid phenol extraction method as described in detail in [38]. RNA concentrations were measured on Nanodrop. The isolated RNA was treated with DNase (Invitrogen AM2238, Turbo DNase) for genomic DNA removal and used for cDNA synthesis. cDNA synthesis was done on equal amounts of DNase treated RNA using PrimeScript 1st strand cDNA synthesis kit and random hexamers (Takara Bio, 6110A) following manufacturer’s instructions. RT-qPCR was performed using the KAPA SYBR FAST qRT-PCR kit (KK4602, KAPA Biosystems). *TAF10* was used as the internal normalization control. Fold change was calculated using the 2^−ΔΔCT method. Statistical significance was determined using Student’s *t*-test.

### Generation of Pho84 overexpression strains

The coding sequence of Pho84 along with a FLAG tag sequence at the C terminus was amplified from the genomic DNA using primers designed with restriction overhangs. The amplified fragment was purified using QIAquick PCR purification kit. The plasmid used for overexpression was the p417 expression system with a strong TEF1 promotor. The plasmid was first digested using EcoR1-HF (NEB R3101S), following manufacturer’s instructions. The amplified fragment was then assembled into the digested plasmid by Gibson assembly (NEB E2611S) following molar ratios and protocols described. The assembled plasmid was used to transform competent *E.coli* DH5α. Plasmids were isolated from obtained clones and they were confirmed for correct ligation by PCR and further by Sanger sequencing. The positive plasmids were then used to transform yeast and positive transformants were confirmed by western blotting for Pho84 expression.

### ATP estimation

ATP levels were measured by ATP estimation kit (Thermo Fisher A22066) as described in [39]. The required strains were grown up to an OD600 of ∼0.8–1.0 (log phase), and cells corresponding to 10 OD600 were pelleted by centrifugation at 1000 × *g* for 2 min at 4 °C. The pellet was resuspended in 300 µl of 5% TCA and incubated on ice for 15 min. The suspension was diluted using 20 mM Tris-HCl, pH 7.0, such that the final TCA concentration was 0.1%; i.e., 6 µl of the sample was diluted with 294 µl of 20 mM Tris-HCl. The reaction mixture for ATP measurement was prepared as per the manufacturer’s instructions, and luminescence was measured using a luminometer. The relative ATP amounts were calculated, and graphs were plotted using GraphPad Prism. Statistical significance was calculated using unpaired Student’s t-test.

### Growth Curve

Required strains were grown in YPD, and cells corresponding to 0.1 OD600 were inoculated into 600 µl YPD in a 48-well plate for growth over 24 h. The experiment was performed with *n* = 3 biological replicates at 30 °C and 200 rpm with continuous orbital shaking. Optical density at 600 nm (OD600) was recorded every 10 min for approximately 24 h using a BioTek multimode plate reader. The growth curves were plotted in R.

### Metabolite extraction and LC-MS/MS analysis

Metabolite extraction and subsequent LC-MS/MS analyses were done using protocols as described in [40,58]. Briefly the required strains were grown in high Pi media to log phase. At an OD600 of ∼0.8-1, equal OD600 cells were quenched in cold 60% methanol (maintained at −45°C) solution for 5 minutes. Cells were then collected by centrifugation (1000xg, 3 min, −5°C), supernatants discarded and cell pellets washed once with quenching buffer. Intracellular metabolites were extracted by first resuspending the pellet in 75% ethanol and heating it at 80°C for 3 minutes followed by ice incubation for ∼1 minute. The extracts were centrifuged at top speed to clear off debris, supernatants containing metabolites were collected and vacuum dried and stored at −80°C until further use.

Metabolites were resuspended in MS grade water and resolved on a Synergi 4-μm Fusion-RP 80 Å (150 × 4.6 mm) LC column (Phenomenex, 00F-4424-E0) using the Shimadzu Nexera UHPLC system. Glycolytic intermediates were separated using two solvent systems; Solvent A: 100% water+ 5 mM ammonium acetate and Solvent B: 100% acetonitrile. Chromatographic flow parameters followed previously established settings [40,58]. Metabolite detection was performed using an AB Sciex Qtrap 5500 mass spectrometer and data was acquired using Analyst 1.6.2 software (Sciex). All glycolytic intermediates were detected on a negative polarity mode. Parent and fragment ion masses for each metabolite are described in [40]. Quantification was carried out by calculating peak areas using MultiQuant software (version 3.0.1).

### Phylogenetic Distribution of Fungal Phosphate Transporters

To establish the underlying evolutionary framework, a reference species tree was constructed using an orthologous gene-based approach based on 25 conserved eukaryotic genes of bacterial ancestry (euBac) [59]. Briefly, homologous sequences were identified and extracted using profile HMMs via HMMER v3.3.2 [60]. Sequences were aligned using MUSCLE v5 [61], and phylogenetically informative regions were retained using BMGE [62]. A concatenated maximum likelihood species tree was inferred using IQ-TREE, utilizing partitioned models specific to each gene (ModelFinder -m TEST option of IQTREE) and evaluated with 1,000 ultrafast bootstrap approximations, with an archaeon *Halobaculum salinum* as the outgroup. The resulting tree was visualized and specifically pruned to the 193 fungal taxa using the iTOL server [63]. This phylogenetic tree and the identification and mapping of high-affinity (PHO84, PHO89) and low-affinity (Pho87, Pho90) phosphate transporters onto this pruned fungal framework were performed as previously described [41]

### Gene Tree Construction and Phylogenetic Analysis

To resolve the evolutionary relationships between the distinct phosphate transporter families, a maximum likelihood gene tree was constructed. Protein sequences comprising fungal PHO84, fungal PHO89, fungal low-affinity transporters, and plant PHT1 were retrieved and subjected to multiple sequence alignment using MUSCLE v5. The resulting alignment was used to infer a maximum likelihood phylogeny using IQ-TREE. The most appropriate sequence evolution model was selected automatically using ModelFinder (-m TEST), and branch support was evaluated using 1,000 ultrafast bootstrap replicates.

## Acknowledgements

We acknowledge the extensive instrument use at the NCBS-TIFR and BRIC inStem mass spectrometry facility. S.L. acknowledges funding support from DBT-Wellcome Trust India Alliance Senior Fellowship (IA/S/21/2/505922) for this study, and support from the Dept. of Biotechnology, Govt. of India (grant IC-12025(22)/4/2023-ICD-DBT) and the DBT-S. Ramachandran National Bioscience Award for Career Development.

## Author contributions

SS and SL conceptualized this study, SS, KS lead experimental design, SS, KS, EM, FC performed experiments, SS, KS, GS and SL analysed data, SS and SL wrote the manuscript. All authors edited the manuscript.

## References

1 Westheimer FH (1987) Why nature chose phosphates. Science (1979) 235, 1173–1178.

2 Hunter T (2012) Why nature chose phosphate to modify proteins. Philosophical Transactions of the Royal Society B: Biological Sciences 367, 2513–2516.

3 Kamerlin SCL, Sharma PK, Prasad RB & Warshel A (2013) Why nature really chose phosphate. Q Rev Biophys 46, 1–132.

4 Van Heerden JH, Wortel MT, Bruggeman FJ, Heijnen JJ, Bollen YJM, Planqué R, Hulshof J, O’Toole TG, Wahl SA & Teusink B (2014) Lost in transition: Start-up of glycolysis yields subpopulations of nongrowing cells. Science (1979) 343.

5 Gupta R & Laxman S (2021) Cycles, sources, and sinks: Conceptualizing how phosphate balance modulates carbon flux using yeast metabolic networks. Elife 10, 1–14.

6 Nicholls JWF, Chin JP, Williams TA, Lenton TM, O’Flaherty V & McGrath JW (2023) On the potential roles of phosphorus in the early evolution of energy metabolism. Front Microbiol 14, 1239189.

7 Wagner CA (2023) The basics of phosphate metabolism. Nephrology Dialysis Transplantation 39, 190.

8 Austin S & Mayer A (2020) Phosphate Homeostasis − A Vital Metabolic Equilibrium Maintained Through the INPHORS Signaling Pathway. Front Microbiol 11, 1–21.

9 Mouillon JM & Persson BL (2006) New aspects on phosphate sensing and signalling in Saccharomyces cerevisiae. FEMS Yeast Res 6, 171–176.

10 Ogawa N, DeRisi J & Brown PO (2000) New components of a system for phosphate accumulation and polyphosphate metabolism in Saccharomyces cerevisiae revealed by genomic expression analysis. Mol Biol Cell 11, 4309–4321.

11 Secco D, Wang C, Shou H & Whelan J (2012) Phosphate homeostasis in the yeast Saccharomyces cerevisiae, the key role of the SPX domain-containing proteins. FEBS Lett 586, 289–295.

12 Lenburg ME & O’Shea EK (1996) Signaling phosphate starvation. Trends Biochem Sci 21, 383–387.

13 Kaffman A, Rank NM & O’Shea EK (1998) Phosphorylation regulates association of the transcription factor Pho4 with its import receptor Pse1/Kap121. Genes Dev 12, 2673–2683.

14 O’Neill EM, Kaffman A, Jolly ER & O’Shea EK (1996) Regulation of PHO4 nuclear localization by the PHO80-PHO85 cyclin-CDK complex. Science 271, 209–212.

15 Kaffman A, Rank NM, O’Neill EM, Huang LS & O’Shea EK (1998) The receptor Msn5 exports the phosphorylated transcription factor Pho4 out of the nucleus. Nature 1998 396:6710 396, 482–486.

16 Komeili A & O’Shea EK (1999) Roles of Phosphorylation Sites in Regulating Activity of the Transcription Factor Pho4. Science (1979) 284, 977–980.

17 Bun-Ya M, Nishimura M, Harashima S & Oshima Y (1991) The PHO84 gene of Saccharomyces cerevisiae encodes an inorganic phosphate transporter. Mol Cell Biol 11, 3229–3238.

18 Lemire JM, Willcocks T, Halvorson HO & Bostian KA (1985) Regulation of repressible acid phosphatase gene transcription in Saccharomyces cerevisiae. Mol Cell Biol 5, 2131–2141.

19 Persson BL, Lagerstedt JO, Pratt JR, Pattison-Granberg J, Lundh K, Shokrollahzadeh S & Lundh F (2003) Regulation of phosphate acquisition in Saccharomyces cerevisiae. Curr Genet 43, 225–244.

20 Levy S, Kafri M, Carmi M & Barkai N (2011) The competitive advantage of a dual-transporter system. Science 334, 1408.

21 Auesukaree C, Homma T, Kaneko Y & Harashima S (2003) Transcriptional regulation of phosphate-responsive genes in low-affinity phosphate-transporter-defective mutants in Saccharomyces cerevisiae. Biochem Biophys Res Commun 306, 843–850.

22 Wykoff DD & O’Shea EK (2001) Phosphate Transport and Sensing in Saccharomyces cerevisiae. Genetics 159, 1491–1499.

23 Hürlimann HC, Pinson B, Stadler-Waibel M, Zeeman SC & Freimoser FM (2009) The SPX domain of the yeast low-affinity phosphate transporter Pho90 regulates transport activity. EMBO Rep 10, 1003–1008.

24 Tamai Y, Toh-e A & Oshima Y (1985) Regulation of inorganic phosphate transport systems in Saccharomyces cerevisiae. J Bacteriol 164, 964–968.

25 Martinez P & Persson BL (1998) Identification, cloning and characterization of a derepressible Na+-coupled phosphate transporter in Saccharomyces cerevisiae. Molecular and General Genetics 258, 628–638.

26 Ribeiro-Filho N, Linforth R, Bora N, Powell CD & Fisk ID (2022) The role of inorganic-phosphate, potassium and magnesium in yeast-flavour formation. Food Research International 162, 112044.

27 Martín JF (2004) Phosphate control of the biosynthesis of antibiotics and other secondary metabolites is mediated by the PhoR-PhoP system: An unfinished story. J Bacteriol 186, 5197–5201.

28 Carstensen A, Herdean A, Schmidt SB, Sharma A, Spetea C, Pribil M & Husted S (2018) The Impacts of Phosphorus Deficiency on the Photosynthetic Electron Transport Chain. Plant Physiol 177, 271.

29 Bechtaoui N, Rabiu MK, Raklami A, Oufdou K, Hafidi M & Jemo M (2021) Phosphate-Dependent Regulation of Growth and Stresses Management in Plants. Front Plant Sci 12, 679916.

30 Khan F, Siddique AB, Shabala S, Zhou M & Zhao C (2023) Phosphorus Plays Key Roles in Regulating Plants’ Physiological Responses to Abiotic Stresses. Plants 12, 2861.

31 Plaxton WC & Tran HT (2011) Metabolic Adaptations of Phosphate-Starved Plants. Plant Physiol 156, 1006–1015.

32 Ceasar SA, Baker A & Ignacimuthu S (2017) Functional characterization of the PHT1 family transporters of foxtail millet with development of a novel Agrobacterium-mediated transformation procedure. Scientific Reports 2017 7:1 7, 14064-.

33 Mitsukawa N, Okumura S, Shirano Y, Sato S, Kato T, Harashima S & Shibata D (1997) Overexpression of an Arabidopsis thaliana high-affinity phosphate transporter gene in tobacco cultured cells enhances cell growth under phosphate-limited conditions. Proc Natl Acad Sci U S A 94, 7098–7102.

34 Nussaume L, Kanno S, Javot H, Marin E, Pochon N, Ayadi A, Nakanishi TM & Thibaud MC (2011) Phosphate Import in Plants: Focus on the PHT1 Transporters. Front Plant Sci 2, 83.

35 Auesukaree C, Homma T, Tochio H, Shirakawa M, Kaneko Y & Harashima S (2004) Intracellular Phosphate Serves as a Signal for the Regulation of the PHO Pathway in Saccharomyces cerevisiae. Journal of Biological Chemistry 279, 17289–17294.

36 Van Dijken JP, Bauer J, Brambilla L, Duboc P, Francois JM, Gancedo C, Giuseppin MLF, Heijnen JJ, Hoare M, Lange HC, Madden EA, Niederberger P, Nielsen J, Parrou JL, Petit T, Porro D, Reuss M, Van Riel N, Rizzi M, Steensma HY, Verrips CT, Vindeløv J & Pronk JT (2000) An interlaboratory comparison of physiological and genetic properties of four Saccharomyces cerevisiae strains. Enzyme Microb Technol 26, 706–714.

37 Sengottaiyan P, Ruiz-Pavõn L & Persson BL (2013) Functional expression, purification and reconstitution of the recombinant phosphate transporter Pho89 of Saccharomyces cerevisiae. FEBS Journal 280, 965–975.

38 Gupta R, Walvekar AS, Liang S, Rashida Z, Shah P & Laxman S (2019) A tRNA modification balances carbon and nitrogen metabolism by regulating phosphate homeostasis. Elife 8.

39 Vengayil V, Niphadkar S, Adhikary S, Varahan S & Laxman S (2024) The deubiquitinase Ubp3/Usp10 constrains glucose-mediated mitochondrial repression via phosphate budgeting. Elife 12.

40 Niphadkar S, Sreedharan S, Vengayil V & Laxman S (2025) Protocol to quantitatively assess glycolysis and related carbon metabolic fluxes using stable isotope tracing in Crabtree-positive yeasts. STAR Protoc 6, 103786.

41 Muthu G, Laxman S, Sai A & Seshasayee N (2025) Evolutionary analysis of Trehalose breakdown pathways. bioRxiv, 2025.10.29.685264.

42 Poirier Y, Jaskolowski A & Clúa J (2022) Phosphate acquisition and metabolism in plants. Current Biology 32, R623–R629.

43 Błaszczyk JW (2023) Metabolites of Life: Phosphate. Metabolites 13, 860.

44 Bowler MW, Cliff MJ, Waltho JP & Blackburn GM (2010) Why did Nature select phosphate for its dominant roles in biology? New Journal of Chemistry 34, 784–794.

45 Zhang Z, Liao H & Lucas WJ (2014) Molecular mechanisms underlying phosphate sensing, signaling, and adaptation in plants. J Integr Plant Biol 56, 192–220.

46 Lamarche MG, Wanner BL, Crépin S & Harel J (2008) The phosphate regulon and bacterial virulence: A regulatory network connecting phosphate homeostasis and pathogenesis. FEMS Microbiol Rev 32, 461–473.

47 Bergwitz C & Jüppner H (2011) Phosphate sensing. Adv Chronic Kidney Dis 18, 132.

48 Bru S, Martínez-Laínez JM, Hernández-Ortega S, Quandt E, Torres-Torronteras J, Martí R, Canadell D, Ariño J, Sharma S, Jiménez J & Clotet J (2016) Polyphosphate is involved in cell cycle progression and genomic stability in *Saccharomyces cerevisiae*. Mol Microbiol 101, 367–380.

49 Bru S, Mayer LM, Kim G-D, Qiu D, Jessen HJ & Mayer A (2025) Acidocalcisome-like vacuoles constitute a feedback-controlled phosphate buffering system for the cytosol. Elife 14.

50 Wang Y, Wang F, Lu H, Liu Y & Mao C (2021) Phosphate Uptake and Transport in Plants: An Elaborate Regulatory System. Plant Cell Physiol 62, 564–572.

51 Popova Y, Thayumanavan P, Lonati E, Agrochão M & Thevelein JM (2010) Transport and signaling through the phosphate-binding site of the yeast Pho84 phosphate transceptor. Proceedings of the National Academy of Sciences 107, 2890–2895.

52 Yip HM, Cheng S, Olson EJ, Crone M & Maerkl SJ (2023) Perfect adaptation achieved by transport limitations governs the inorganic phosphate response in S. cerevisiae. Proc Natl Acad Sci U S A 120, e2212151120.

53 Springer M, Wykoff DD, Miller N & O’Shea EK (2003) Partially Phosphorylated Pho4 Activates Transcription of a Subset of Phosphate-Responsive Genes. PLoS Biol 1, e28.

54 Wykoff DD, Rizvi AH, Raser JM, Margolin B & O’Shea EK (2007) Positive Feedback Regulates Switching of Phosphate Transporters in S. cerevisiae. Mol Cell 27, 1005–1013.

55 Wykoff DD & O’Shea EK (2001) Phosphate transport and sensing in Saccharomyces cerevisiae. Genetics 159, 1491.

56 Longtine MS, McKenzie A, Demarini DJ, Shah NG, Wach A, Brachat A, Philippsen P & Pringle JR (1998) Additional modules for versatile and economical PCR-based gene deletion and modification in Saccharomyces cerevisiae. Yeast 14, 953–961.

57 Niphadkar S, Karinje L & Laxman S (2024) The PP2A-like phosphatase Ppg1 mediates assembly of the Far complex to balance gluconeogenic outputs and enables adaptation to glucose depletion. PLoS Genet 20, e1011202.

58 Walvekar A, Rashida Z, Maddali H & Laxman S (2018) A versatile LC-MS/MS approach for comprehensive, quantitative analysis of central metabolic pathways [version 1; referees: 2 approved]. Wellcome Open Res 3, 1–15.

59 He D, Fiz-Palacios O, Fu CJ, Tsai CC & Baldauf SL (2014) An alternative root for the eukaryote tree of life. Current Biology 24, 465–470.

60 Eddy SR (2011) Accelerated Profile HMM Searches. PLoS Comput Biol 7, e1002195.

61 Edgar RC (2022) Muscle5: High-accuracy alignment ensembles enable unbiased assessments of sequence homology and phylogeny. Nature Communications 2022 13:1 13, 6968-.

62 Criscuolo A & Gribaldo S (2010) BMGE (Block Mapping and Gathering with Entropy): a new software for selection of phylogenetic informative regions from multiple sequence alignments. BMC Evolutionary Biology 2010 10:1 10, 210-.

63 Letunic I & Bork P (2021) Interactive Tree Of Life (iTOL) v5: an online tool for phylogenetic tree display and annotation. Nucleic Acids Res 49, W293–W296.

